# Inhibition of adhesive nanofibrillar mediated *Streptococcus gordonii - Candida albicans* mono- and dual-species biofilms

**DOI:** 10.1101/689661

**Authors:** Raja Veerapandian, Govindsamy Vediyappan

## Abstract

Dental caries and periodontitis are the most common oral disease of all age groups, affecting billions of people worldwide. These oral diseases are mostly associated with the microbial biofilms in the oral cavity. *Streptococcus gordonii*, an early tooth colonizing bacterium and *Candida albicans*, an opportunistic pathogenic fungus, are the two abundant oral microbes form mixed biofilms and augment their virulence properties affecting oral health negatively. Understanding the molecular mechanisms of their interactions and blocking the growth of these biofilms by nontoxic compounds could help develop effective therapeutic approaches. We report in this study, inhibition of mono- or dual-species biofilms of *S. gordonii* and *C. albicans*, and biofilm eDNA *in vitro* by Gymnemic Acids (GAs), a nontoxic small molecule inhibitor of fungal hyphae. Scanning electron microscopic images of biofilms revealed attachment of *S. gordonii* cells to the hyphal and on saliva-coated hydroxyapatite (sHA) surfaces via nanofibrils only in the untreated control but not in the GAs treated biofilms. Interestingly, *C. albicans* produced fibrillar adhesive structures from hyphae when grown with *S. gordonii* as mixed biofilm and addition of GAs to this biofilm abrogates the nanofibrils, reduces the growth of hyphae, and biofilms. To our knowledge, this is a first report that *C. albicans* produces adhesive fibrils from hyphae in response to *S. gordonii* mixed biofilm growth. A semi-quantitative PCR data of selected genes related to biofilms of both microbes show their differential expression. Further evaluation of one of the gene products of *S. gordonii* revealed that GAs could inhibit its recombinant glyceraldehyde-3-phosphate dehydrogenase (GAPDH) enzyme activity. Taken together, our results suggest that *S. gordonii* stimulates expression of adhesive materials in *C. albicans* by direct interaction and or by signaling mechanism(s), and these mechanisms can be inhibited by GAs. Further studies on global gene expression of these biofilms and their biochemical studies may reveal the molecular mechanism of their inhibition.

## Introduction

Dental caries is a polymicrobial biofilm-induced disease affecting 3.5 billion people globally (Kassebaum et al., 2017). The worldwide annual total costs due to dental diseases are estimated around $545 billions in 2015 (Righolt et al., 2018). *Candida albicans* fungus is the etiologic agent of oral thrush and denture stomatitis, the two mucosal oral biofilm infections in immunocompromised patients and in aged peoples, respectively. *C. albicans* and *Streptococcus* bacterial species are abundant in the oral cavity and readily form mixed biofilms which are resistant to antimicrobials and serves as a source for systemic infections (Dongari-Bagtzoglou et al., 2009;Silverman et al., 2010;Diaz et al., 2012;Ricker et al., 2014;O’Donnell et al., 2015). Some of the streptococci (e.g. *S. mutans*) are the causative agents of dental caries and gum disease. Recent studies have shown that a complex interaction and aggregation occurs between streptococci and *C. albicans*, and the molecular mechanisms are poorly understood (Dutton et al., 2014;Hwang et al., 2017).

*C. albicans* is a commensal and an opportunistic human fungal pathogen found in cutaneous, oral, intestine, genital regions, and can initiate various forms of Candidiasis. While *C. albicans* is the primary causative agent of oral thrush or oropharyngeal candidiasis in the immune compromised populations (Odds, 1987), various groups of oral bacteria are shown to interact with *C. albicans* and influence the disease severity (Dongari-Bagtzoglou et al., 2009;Harriott and Noverr, 2011). Oral streptococcal species including *Streptococcus gordonii, S. oralis*, and *S. mutans* interact with *C. albicans* and augment both fungal and bacterial virulence (Silverman et al., 2010;Ricker et al., 2014;O’Donnell et al., 2015;Hwang et al., 2017). Other bacteria including *Staphylococcus aureus* (Harriott and Noverr, 2009) and *Acinetobacter baumannii* (Uppuluri et al., 2018) use *C. albicans* hyphae as a substratum for attachment, and forms robust biofilms.

*C. albicans* exists in yeast, pseudohyphae and hyphae. The transition from yeast or pseudohyphae to hyphae is required for its tissue invasion and biofilm formation. Mutants that are defective in hyphal growth are avirulent and unable to form biofilms (Lo et al., 1997;Nobile and Mitchell, 2006). Hence, *C. albicans* hyphae play a pivotal role in biofilms growth and virulence. Some of the oral bacteria, including *S. gordonii* are shown to promote the hyphal growths of *C. albicans* and bind preferably to these hyphal surfaces (Bamford et al., 2009). This hyphal binding increased the biofilm mass, and chemical inhibition of candida hyphae reduced the biofilm mass (Bamford et al., 2009). Several bacterial pathogens exploit *C. albicans* hyphae for their attachments (Silverman et al., 2010;Diaz et al., 2012;Dutton et al., 2014;Xu et al., 2014b;O’Donnell et al., 2015). A recent study has shown that the yeast cells of *C. glabrata* bind to *C. albicans* hyphae and forms fungal-fungal biofilms in the oral milieu (Tati et al., 2016). Mono- and mixed biofilms are highly resistant to antimicrobial agents and serve as reservoirs for systemic dissemination, cause inflammation (Nett et al., 2010;Vediyappan et al., 2010;Xu et al., 2014b) and can sequester antimicrobial drugs (Nett et al., 2010;Vediyappan et al., 2010). It is plausible that inhibiting *C. albicans* hyphal growth by nontoxic small molecules could abrogate the hyphae related virulence, including its co-interaction with bacteria and growth of polymicrobial biofilms.

Gymnemic acids (GAs), a family of triterpenoid molecules from *Gymnema sylvestre* medicinal plant was shown to block *C. albicans* yeast to hypha transition and hyphal growth *in vitro* and in a worm *(Caenorhabditis elegans)* model of invasive candidiasis(Vediyappan et al., 2013). GAs contain various pharmacological properties including antagonistic activity against the ß-isoform of Liver-X-Receptor (LXR) which could result decreased lipid accumulation in liver cells (Renga et al., 2015), modifying sweet taste sensation by binding to taste receptors, T1R2, and T1R3 (Sanematsu et al., 2014), and blocking the uptake of glucose in the intestinal cells (Wang et al., 2014). The GAs rich gymnema extract has been used in humans for treating diabetes and obesity (Baskaran et al., 1990;Porchezhian and Dobriyal, 2003;Leach, 2007). A recent clinical study confirmed the traditional use of *G. sylvestre* for diabetes (Zuniga et al., 2017). We have shown that GAs inhibit the growth of hyphae (polarized growth) without affecting the yeast form of growth (isometric growth) (Vediyappan et. al. 2013) and GAs are likely acting via hyphal growth regulatory pathways (unpublished results). Since GAs block the hyphal growth of *C. albicans* and *S. gordonii* or other oral bacteria use *C. albicans* hyphae for their attachment (Bamford et al., 2009) and for mixed biofilm growths, we wanted to test the hypothesis that preventing *C. albicans* hyphae by GAs could abolish bacteria-C. *albicans* interactions and their mixed biofilms. In the current study, we show a synergistic interaction between *S. gordonii* and *C. albicans in vitro* and the addition of gymnemic acids (GAs) prevented the growth of mono- or dual-species biofilms. Our results show, for the first time to our knowledge, formation of ‘nanofibrillar’ structures from *C. albicans* hyphae in response to *S. gordonii* co-culture, which correlates their enhanced interaction and biofilms growth. Treating mono- or dual-species biofilms with GAs abolished these structures and reduced their biofilm growths.

## Materials and Methods

### Strains and culture conditions

*Streptococcus gordonii* ATCC 10558 (generously provided by Dr. Indranil Biswas, Kansas University Medical Centre, KU, Kansas City) and *Candida albicans* SC5314 (genome sequenced) were used to generate mono- or dual-species biofilms in 24-well microtiter plates (Costar) under static condition. *E. coli* 10-Beta (NEB) and BL21(DE3) (Novagen, Madison, WI) were used for cloning and expression of recombinant proteins and were routinely grown in Luria-Bertani (LB) broth or on LB agar. GAs was purified and evaluated for their HPLC profiles according to the published protocols (Vediyappan et al., 2013;Sanematsu et al., 2014) and mixture of GAs was used in this study. Minimum biofilm inhibitory concentration (MBIC) was determined in 24-well plate as previously described (Saputo et al., 2018) with slight modifications using TYES broth medium (1% tryptone and 0.5% yeast extract at pH 7.0 with 1% (wt/vol) sucrose). The MBIC is defined as the lowest concentration of GAs that inhibit maximum amount of biofilm growth. Briefly, the suspension of *S. gordonii* was added to a 24-well plate containing a serially diluted GAs at concentrations ranging from 0 to 1000 μg/mL and incubated at 37 °C with 5% CO2 for 18 h. Medium with and without GAs served as controls. After washing off unbound cells and medium with PBS, the adhered biofilms were measured by crystal violet (0.1%, CV) staining (Merritt et al., 2005). Experiments were repeated at least three times each with triplicates, and representative results are shown.

To determine the effect of GAs on the growth rate of *S. gordonii*, we used a Bioscreen-C real time growth monitoring system (Oy Growth Curves Ab Ltd, Finland). In this method, 200 μl of growth medium containing exponentially growing *S. gordonii* cells were added into the honeycomb wells (triplicate) with or without GAs (control). Wells with different concentrations of GAs (in 200 μl total volume) were served as GAs treated. The plate was incubated at 37 ^o^ C without shaking except a 10-second shaking before reading absorbance at 600 nm at 30-minute intervals. The overall objective of the kinetic growth reading of *S. gordonii* in the presence or absence of GAs was to determine if GAs exert toxic effect on *S. gordonii* cells.

### Unstimulated whole saliva preparation

Human saliva collection and processing were done as described previously (Jack et al., 2015). Briefly, unstimulated whole human saliva was collected from at least 5-6 healthy volunteers with Institutional Review Board (IRB) protocol approval (#9130.1) from Kansas State University. All the subjects gave written informed consent approved by the IRB committee. Saliva was pooled and mixed with 2.5 mM dithiothreitol and kept in ice for 10 min before clarification by centrifugation (10,000 × g for 10 min). The supernatant was diluted to 10% in distilled water and filter sterilized through a 0.22-μm nitrocellulose filter and stored at −80°C in aliquots. Diluted saliva was used to coat the microtiter wells and hydroxyapatite (HA) discs (Clarkson Chromatography Products, PA) overnight.

### Mono- and dual-species biofilm assay

To test the effect of GAs on biofilm formation in saliva-coated wells, *S. gordonii* and *C. albicans* were grown alone or in combinations in TYES medium with or without GAs (500 μg/ml) for 18 hours statically at 37°C and 5% CO2. Biofilms of *S. gordonii* and *C. albicans*, either as mono- or as mixed species, were developed for up to 18 h as reported (Dutton et al., 2014;Ricker et al., 2014) except that saliva-coated hydroxyapatite (sHA) discs were also used in the current study. Briefly, sHA discs were placed in a 24-well plate and inoculated with approximately 2 × 10^6^ (CFU/ml) of *S. gordonii* or/ and 2 × 10^4^ (CFU/ml) of *C. albicans* in the TYES medium with or without GAs. The effect of GAs against biofilm formation was studied using CV staining (Merritt et al., 2005). Briefly, biofilm was stained with CV, washed twice with PBS to remove excess dye and solubilized by 95% ethanol. The absorbance of the solutions was read at OD595 by a Victor 3v multimode reader (Perkin Elmer, USA). Experiments were repeated at least three times each with triplicates, and representative results are shown.

### Measurement of biofilm extracellular DNA (eDNA)

eDNA were measured as per the standard protocol described elsewhere (Jack et al., 2015). Briefly, biofilms were scraped from saliva coated wells or sHA disc into 0.5 mL TE buffer (10 mM Tris/HCl, pH 7.5 and 1mM EDTA), cell free DNA was collected by centrifugation at 10,000 × g for 5 min. The DNA concentration was then analyzed from the supernatant using a NanoDrop 2000 spectrophotometer (Thermo Scientific, USA).

### Scanning electron microscopy (SEM)

SEM was done as per the standard protocol described previously (Erlandsen et al., 2004). Briefly, sHA disc with a biofilm on their surface were fixed with 2% paraformaldehyde and 2% glutaraldehyde in 0.15 M sodium cacodylate buffer, pH 7.4, containing 0.15% Alcian blue. Biofilms grown on sHA discs were washed with 0.15 M cacodylate buffer and dehydrated in a graded series of ethanol concentrations. Specimens were mounted on adhesive carbon films and then coated with 1 nm of platinum using an Ion Tech argon ion beam coater. Prepared samples were observed in a SEM (Field Emission Scanning Electron Microscope, Versa 3D Dual Beam, Nikon).

### RNA isolation, cDNA Synthesis and semiquantitative RT–PCR

Biofilm were treated with RNA protect bacteria reagent (Qiagen, Valencia, CA) for 5 min to stabilize RNA and stored at −80°C. Total RNA was isolated from the biofilms using the TRIzol reagent (Invitrogen, Carlsbad, CA) and cDNA was synthesized using the SuperScript III indirect cDNA labelling kit (Invitrogen) as per the manufacturer instructions. The semi-quantitative RT-PCR using 2X PCR Master Mix (Promega Corporation, Madison, WI, USA) and primers was carried out in a 20 μL reaction volume (1 μL cDNA, 10 μL Master Mix, 0.5 μM of each primer). Primer details are given in Tables 1 and 2. The internal control used was *16S rRNA* for *S. gordonii* and *TDH3* for *C. albicans.* The cycling conditions consisted of initial denaturation at 94°C for 3 min followed by denaturation at 94°C for 30 seconds, annealing at 50°C or 58°C for 30 seconds, and extension at 72°C for 45 seconds, then final extension at 72°C for 7 min. Twenty microliters of each PCR product was electrophoresed on agarose gel (1.2% w/v) containing ethidium bromide (0.5 μg/ml). Images of the amplified products were acquired with an Alpha Imager; the intensity was quantified using the image J software (NIH, USA). The band intensity was expressed as mRNA expression in fold (specific gene expression/internal control gene expression). The expression of the control cells without treatment was taken as one and it was compared with the treated group.

**Table 1.**
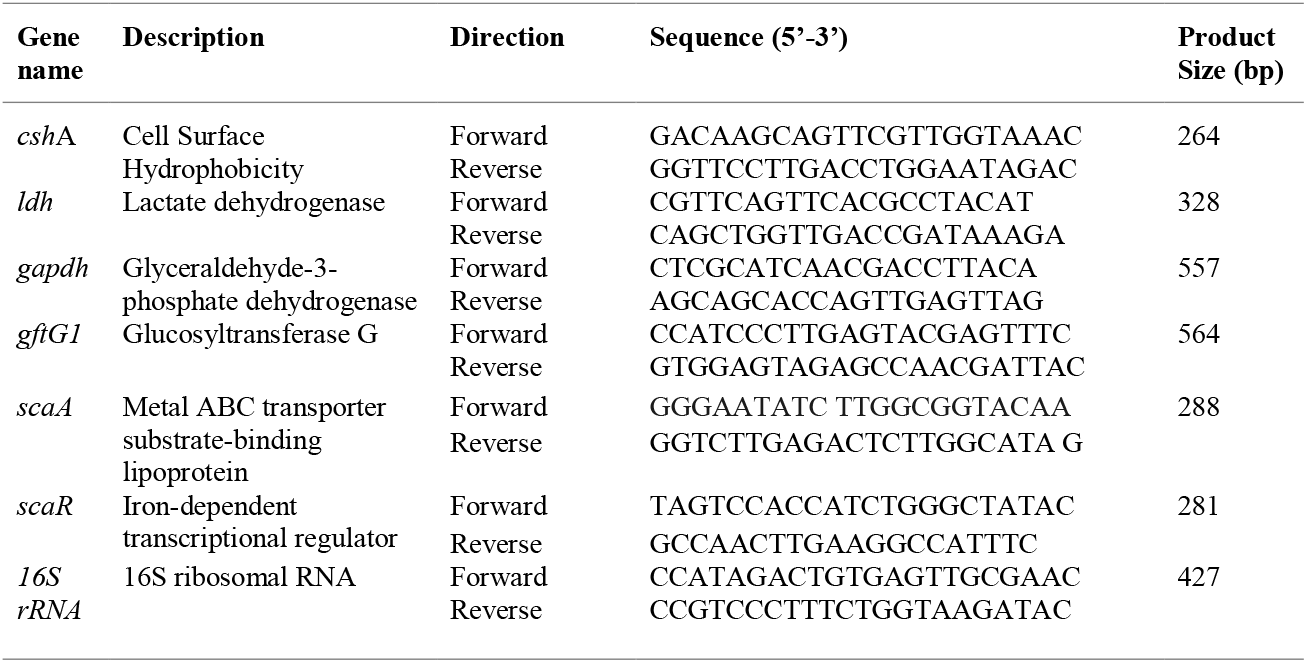
List of *S. gordonii* specific primers used for semi quantitative RT-PCR

**Table 2.**
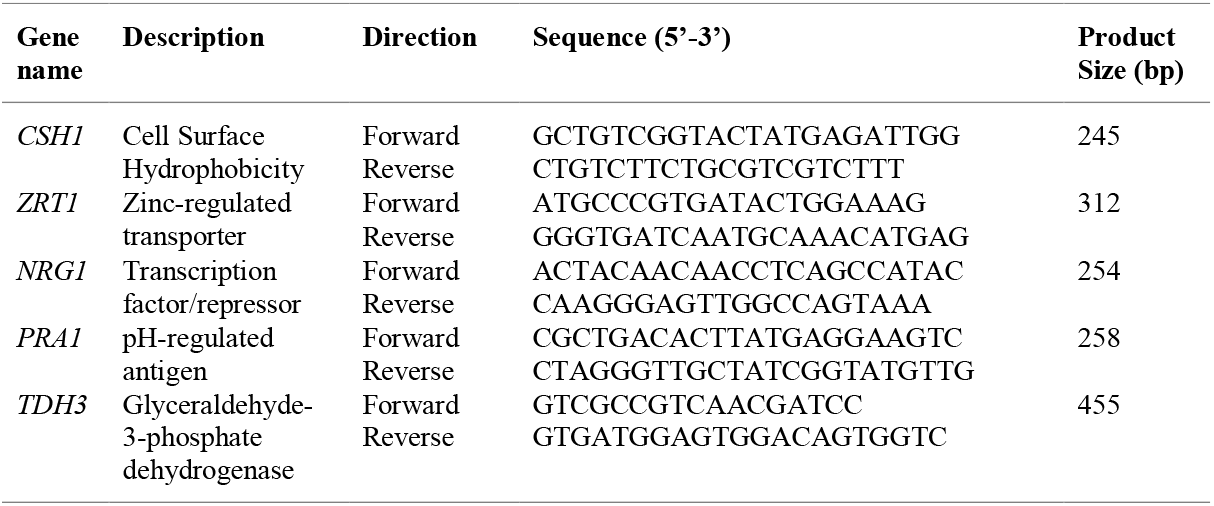
List of *C. albicans* primers used for semi quantitative RT-PCR

### Cloning, Expression, and Purification of rGAPDH

GAPDH gene from *S. gordonii* was PCR amplified using primers GAPDH-F (5′-ATTCCATATGGTAGTTAAAGTTGGTATTAACGGT-3′) and GAPDH-R (5′-GCGCTCGAGTTTAGCGATTTTCGCGAAGTATTCAAG-3′) where the underlined sequences in the forward and in reverse primers indicate NdeI and XhoI restriction sites, respectively. Following PCR amplification of chromosomal DNA from *S. gordonii* strain ATCC 10558, amplicons of 1008 bp were digested with NdeI and XhoI and inserted into the predigested pET28b plasmid (Novagen, Madison, WI). Successful cloning of the gene was confirmed by restriction endonuclease and DNA sequence analyses (data not shown). Recombinant plasmid was transformed into *E. coli* BL21 (DE3) for overexpression. Expression of GAPDH-6His protein was induced with 1mM isopropyl-ß-d-thiogalactopyranoside (IPTG) when cultures reached an optical density at 600 nm (OD600) of 0.6, and cells were harvested after 4 h. The cell pellet from 2 L of culture was resuspended in 40 ml of a buffer containing 50mM NaH2PO4 pH 8.0, 300mM NaCl, 20mM imidazole with 1x protease inhibitor cocktail (Roche) and 1mM phenylmethylsulfonyl fluoride (PMSF), and cells were lysed by French press (~19,000 psi). The lysate was centrifuged at 10,000 g for 20 min at 4°C. Recombinant His-tagged GAPDH was purified using Ni-NTA Agarose (Qiagen, Valencia, USA) in native conditions according to the manufacturer’s recommendations. GAPDH was eluted using gradients of increasing imidazole concentration (100-300mM). Fractions containing rGAPDH were pooled and dialyzed against distilled water and used for subsequent analysis.

### SDS-PAGE and Immunoblotting

The purity of the proteins was checked using SDS-PAGE electrophoresis in a vertical electrophoretic mini-cell unit (Bio-Rad, Hercules, CA), in Tris-glycine running buffer (25mM Tris, 192mM glycine, 0.1% SDS [pH 8.3]), for 1 h at 120 V. Proteins were transferred to Immobilon-P PVDF membrane (pore size, 0.45μm; Millipore Sigma, USA) and blocked with 5% nonfat dry milk in Tris-buffered saline (20mM Tris, 150mM NaCl, 0.2% Tween 20 [pH 7.5]). Membranes were incubated with anti-GAPDH immune sera raised in rabbits, followed by incubation with secondary antibody (anti-rabbit IgG; Cell Signaling Technology, USA). Reacted protein bands were visualized by using Pierce™ ECL 2 Western Blotting Substrate (Thermo scientific, USA) and imaging.

### Determination of GAPDH activity

GAPDH activity was measured in the presence and absence of GAs using Glyceraldehyde 3 Phosphate Dehydrogenase Activity Colorimetric Assay Kit (ab204732, Abcam, Cambridge, MA) as per the manufacturer instructions. Briefly, purified rGAPDH protein (0.1μM) were mixed with and without GAs (100 and 200μM) followed by the addition of reaction mix supplied from the kit components. The conversion of NAD to NADH was monitored for every10s spectrometrically by Victor 3 multimode reader (Perkin Elmer, USA) at 450 nm.

### Statistical analysis

Data from multiple experiments (≥3) were quantified and expressed as Mean± SD, and differences between groups were analyzed by using one-way ANOVA. A *p* ≤ 0.05 was considered significant in all analyses. The data were computed with GraphPad Prism version 7.0 software.

## Results

### Determination of minimum biofilm inhibition concentration (MBIC)

*S. gordonii* and *C. albicans* co-exist in the oral cavity as abundant microbes, and the former is known to attach on the hyphal surfaces of the latter forming a mixed species biofilm with enhanced virulence. Preventing the growth of these biofilms by nontoxic small molecules would limit oral diseases and their dissemination systemically. A previous study from this laboratory showed that GAs prevent *C. albicans* yeast-to-hypha transition and its hyphal growth without affecting the growth of yeast form and viability (Vediyappan et al., 2013). Based on these results, we predicted that lack of hyphae as substrates or its surface molecules due to GAs effect would preclude the binding of bacteria to them and thus could prevent the growth of mixed biofilms. However, the impact of GAs on *S. gordonii*’ biofilms growth and or its hypha inducing effect on *C. albicans* during mixed biofilm growth is unknown. First, we wanted to determine the MBIC of GAs. To determine the minimum amount of GAs needed to inhibit maximum biofilm growth of *S. gordonii*, MBIC assay was performed with increasing concentrations of GAs (0 to 1000 μg/mL). Biofilms were quantified by CV staining and the results showed a concentration-dependent antibiofilm activity of GAs against *S. gordonii* (Figure 1A). A significant inhibition was found from concentration >400 μg/mL.

**Figure 1.**
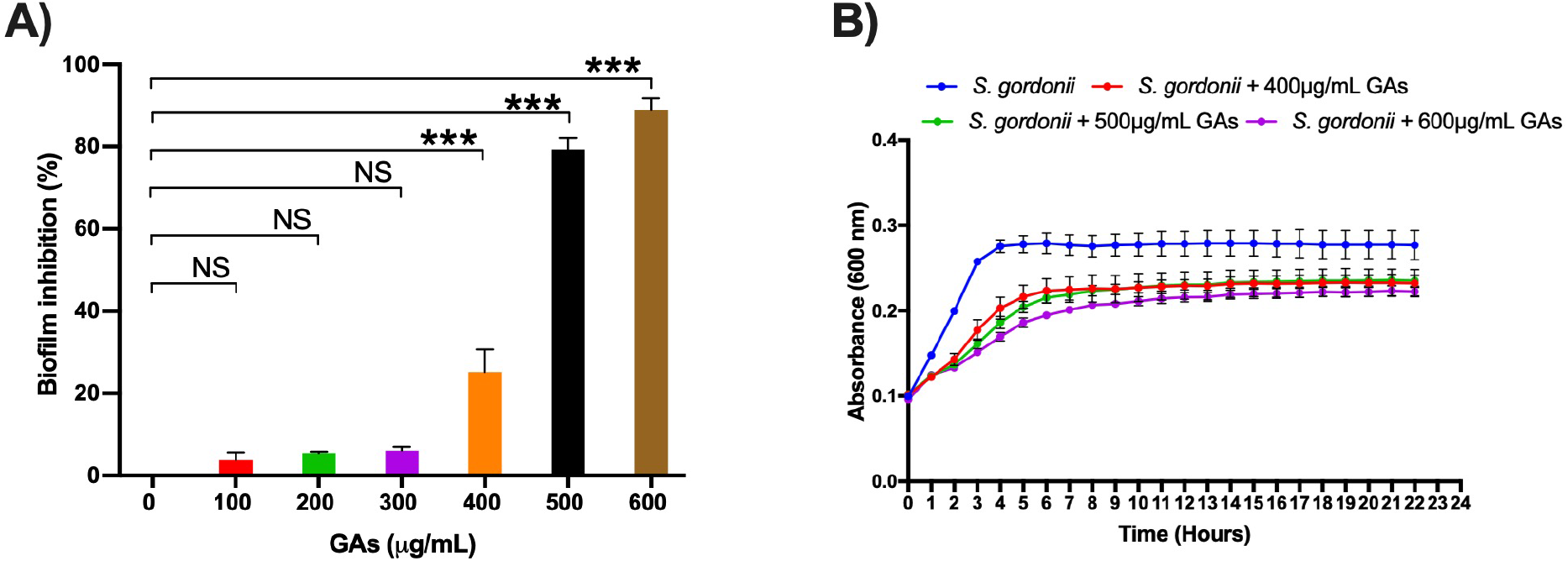
**A) Determination of minimum biofilm inhibition concentration of GAs against *S. gordonii.*** Varying concentrations of GAs (0-1000 μg/ml) were used in TYES medium containing *S. gordonii* in 24-wells with triplicates and incubated at 37 °C with 5% CO2 statically for 18 h. Biofilms grown without GAs served as controls. Inhibition of biofilms growth was analyzed by CV staining and % inhibition of biofilm was calculated. The results represent means ± standard deviations for three independent experiments. Statistical significance was determined by ANOVA. *** *p* < 0.0001, NS = not significant. **B) Analysis of *S. gordonii* planktonic growth with and without GAs**. Effect of GAs at varying concentrations over the planktonic growth of *Streptococcus gordonii.* Honeycomb wells containing *S. gordonii* in 200 μl TYES medium with or without GAs was used to monitor the growth rate for the indicated time. Absorbance was recorded every 30-minute intervals at 600 nm as described in the methods section. The results represent means ± standard deviations for three independent experiments.

To determine if GAs are toxic to *S. gordonii*, we measured its growth kinetics as planktonic cells in the presence or absence of GAs using Bioscreen-C growth monitor at 37 ^o^C in TYES medium as described in the method section. Since GAs inhibited *S. gordonii* biofilms growth from concentration 400 μg/ml or above, we used three different concentrations of GAs (400, 500 & 600 μg/ml) to assess its effects. As shown in Figure 1B, the overall growth kinetics of *S. gordonii* in the presence or absence of GAs is parallel except the wells with GAs showed a reduced growth rate. While *S. gorodonii* exposed to 400 & 500μg/ml GAs show similar growth pattern, GAs at 600μg/ml affect *S. gordonii* growth even further. Taken together, GAs were inhibiting the growth of *S. gordonii* moderately at 400 – 600μg/ml under the growth conditions used. Since 500μg /ml GAs inhibited maximum biofilm growth, we employed this concentration (500μg/ml) throughout the study to determine its effect on mono- or dual-species biofilms.

### Inhibition of *S. gordonii* and *C. albicans* mono- and dual-species biofilms grown in 24-well microtiter plates by GAs

The antibiofilm efficacy of GAs was assessed under *in vitro* condition by measuring the binding of CV to *S. gordonii* biofilms cells in 24-well plates. The antibiofilm activity of GAs was effective at 500μg/ml against *S. gordonii* and *C. albicans* mono- and dual-species biofilms (Figure 2A). GAs treatment significantly reduced the amount of *S. gordonii* biofilms (Figure 2A). Similarly, the mixed biofilms were also reduced with GAs treatment and are significant as analyzed by one-way ANOVA *(p* = 0.001).

**Figure 2.**
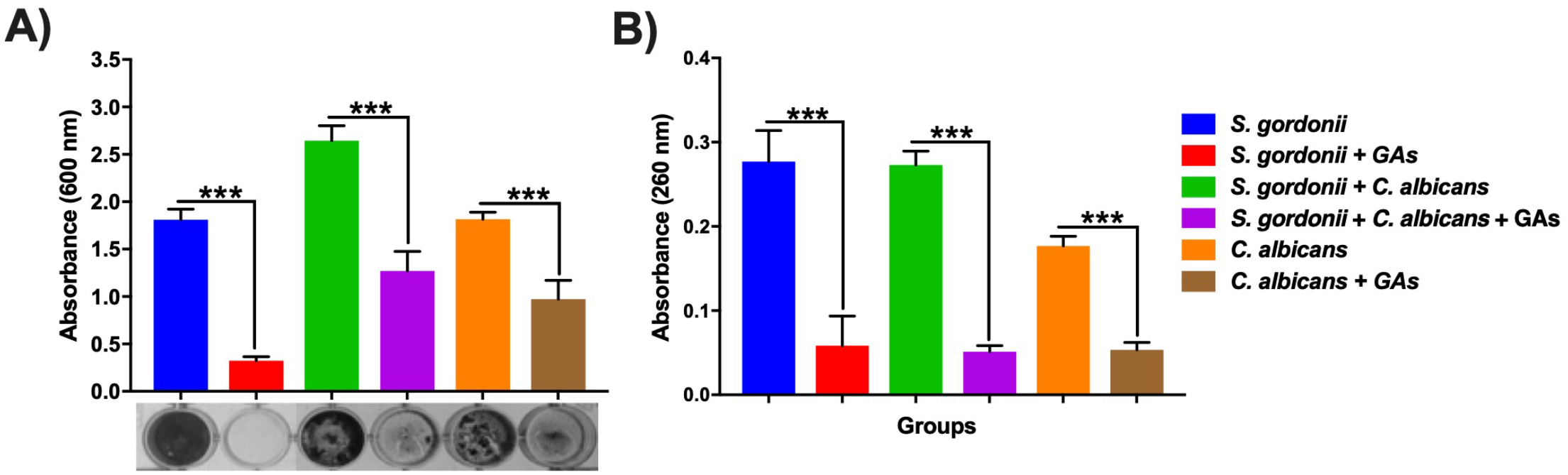
Effect of GAs on *S. gordonii* and *C. albicans* mono- or dual-species biofilms. A) Crystal violet staining of *S. gordonii* and *C. albicans*, either as mono- or as dual-species biofilms with and without GAs in saliva coated 24-well plates. B) Measurement of eDNA from mono- and dual-species biofilms with and without GAs. The results represent means ± standard deviations for three independent experiments. NS-not significant, \**p* < 0.05, \*\**p*<0.01, \*\*\**p* < 0.001.

### GAs treatment effectively reduces eDNA in mono- and dual-species biofilms

Extracellular DNA (eDNA) is known to be released during biofilm growth and is an integral part of polymeric material (Xu and Kreth, 2013). To examine the effects of GAs on biofilm eDNA, mono- and dual-species biofilms were grown in TYES medium. High level of eDNA were found in both mono- and dual-species biofilms. Interestingly, significant reduction in eDNA concentrations were observed in these biofilms treated with GAs *(p* = 0.001, Figure 2B).

### Inhibition of *S. gordonii* and *C. albicans* mono- and dual-species biofilms on sHA discs

SEM analysis was carried out to examine the structures of mono- and dual-species biofilms formed on sHA discs that were treated with and without GAs. Biofilms formed on sHA discs were fixed, stained with alcian blue and processed as described (Erlandsen et al., 2004). SEM micrographs of *S. gordonii* revealed the formation of biofilms with thick aggregates of cells on the surface of sHA containing patches of exopolysaccharide (EPS) (Figure 3A). Interestingly, very little biofilms of *S. gordonii* was found on the GAs treated sHA disc, and large empty areas were seen mostly (Figure 3G). These SEM results agree with *in vitro* biofilm growth assay (Figure 2A). As expected, *C. albicans* control biofilms (B) contain multilayers of hyphae and in the GAs exposed biofilms, very little yeast and pseudohyphal cells were present on the sHA discs (H-I). Although GAs does not inhibit the yeast fungal growth rate at the concentration used (Vediyappan et al., 2013), the yeast-pseudohyphal cells produced due to GAs treatment poorly or could not be attached to the sHA discs or to the other surfaces (unpublished results). *S. gordonii* and *C. albicans* dual biofilms (C) contained both bacterial and fungal hyphal cells densely, and their abundance decreased by GAs treatment (Figure 3I).

**Figure 3.**
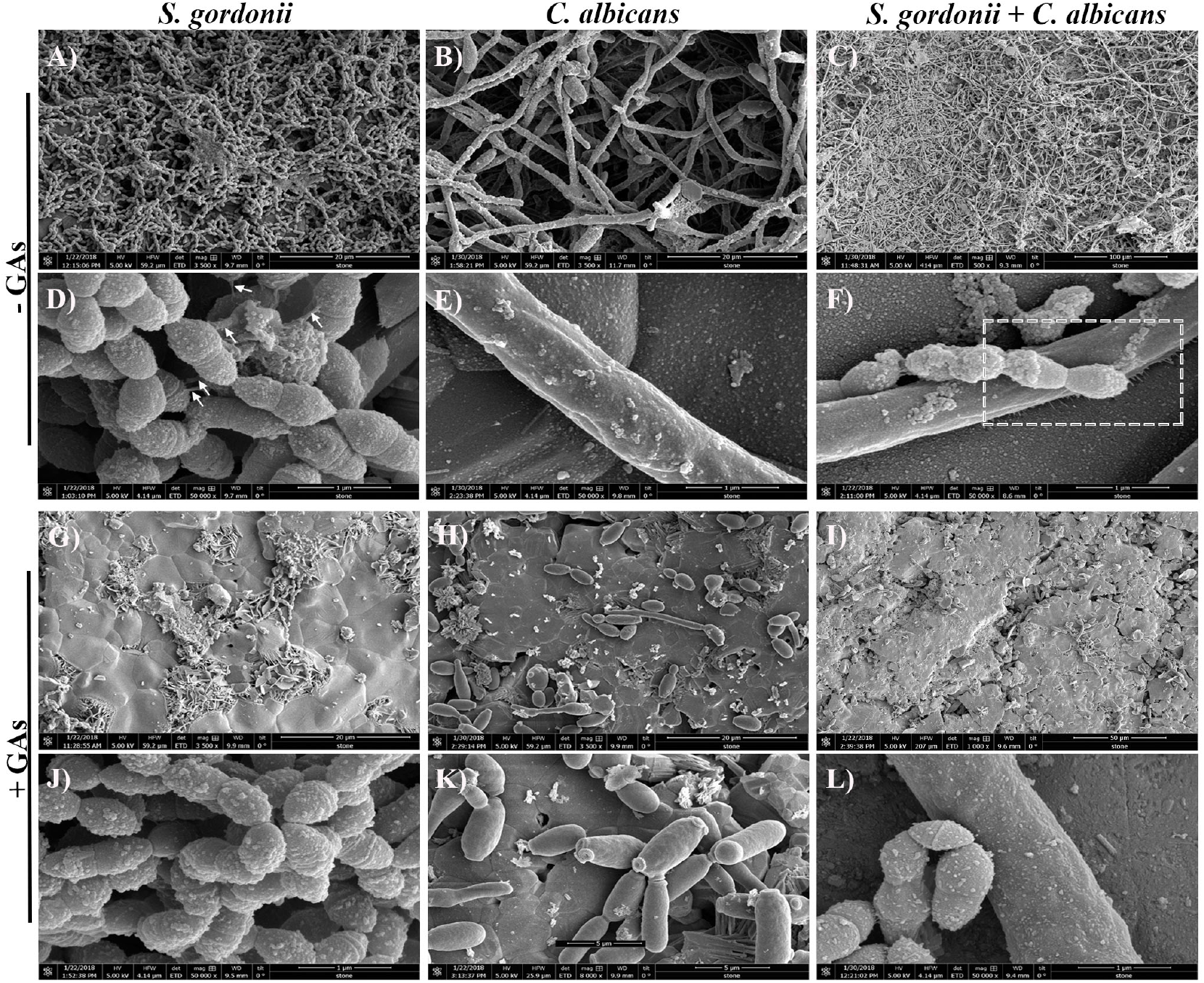
Scanning electron microscopy (SEM) observations of mono- and dual-species biofilms grown on sHA with and without GAs. Images of mono- and dual-species biofilms grown for 18 h in the absence (control, A-F) and in the presence of GAs (treated, G-L) at 500 μg/ml concentration. Biofilms grown with GAs show few cells on the sHA surfaces compared to dense layers of cells with exopolysaccharides (EPS) in the control groups. Short fibrils in the untreated *S. gordonii* biofilms that are attached to neighboring cells are shown (D, arrows). Changes in the biofilm surface textures and absence of fibrils were observed in the GAs treated biofilm groups (G-L). GAs treated *C. albicans* show mostly yeast or pseudohyphal cells with few hyphae (H & K). Dashed box in F was further magnified in figure 4 to show nanofibrillar structures. In GAs treated dual biofilms, weak or no fibrillar structures from *S. gordonii* and none from *C. albicans* were found.

### Treatment with GAs significantly reduced extracellular nanofibrillar based biofilm formation

In order to get better insight, we then viewed these biofilms closely at higher magnification (50, 000x). In *S. gordonii* biofilms without GAs exposure, we found short fibrils between *S. gordonii* cells [and some of these fibrils were attached to sHA] (Figure 3D, arrows). As expected, *C. albicans* biofilms without GAs treatment produced mostly hyphae. Interestingly, *C. albicans* biofilms co-cultured with *S. gordonii* (dual biofilms) without GAs showed several closely attached bacterial-fungal cells with abundant extracellular materials.

Strikingly, we found several thin fibrils from hypha that is in close contact with the sHA disc (Figure 3F, boxed) or to the neighbouring hypha (Figure 4, A-D & arrows). *S. gordonii* exhibits high affinity to the *C. albicans* hyphae as the bacterium coiled around the hypha and also by direct attachment with the help of fibrils. The inhibitory effect of GAs was clearly demonstrated in the SEM micrographs of biofilms. Interestingly, mono- and dual-species biofilms grown on sHA discs treated with GAs were devoid or having less cell surface nanofibrils and exhibited completely smooth hyphal surfaces (Figure 3).

**Figure 4.**
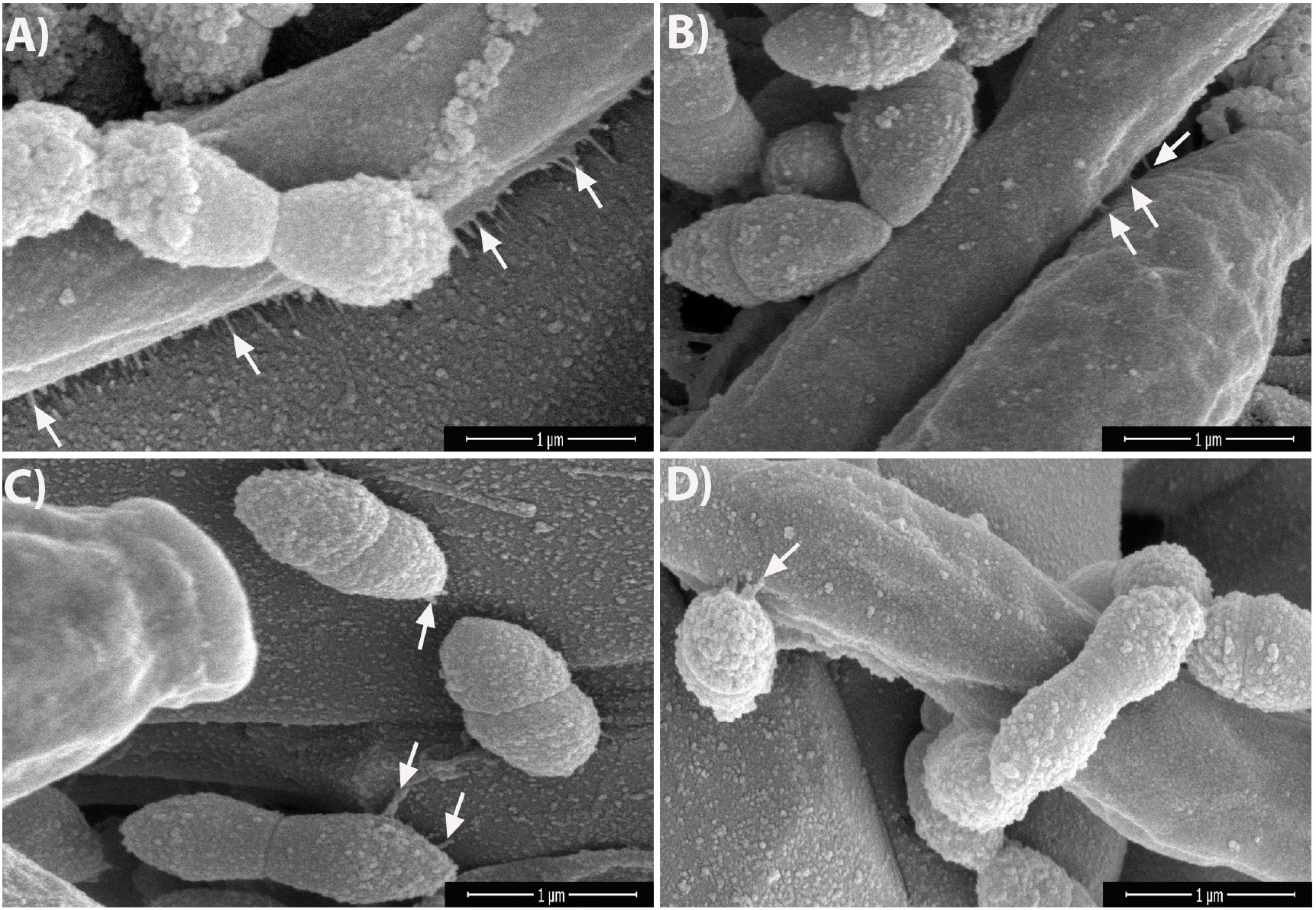
Dual-species biofilms in the absence of GAs promote nanofibrillar-mediated interactions. SEM images of dual-species biofilms showing nanofibrillar structures from *C. albicans* hypha that are attached to the sHA (A, arrows) as well as between two hyphae (B, arrows). *S. gordonii* exhibits high affinity to hypha by its fibrillar attachment and by tight coiling around the hypha (D), and also to sHA (C) which mimics teeth.

### Modulation of gene expression in mono- and dual-species biofilms with and without GAs

Few studies have shown the differential expression of genes during *S. gordonii* (Gilmore et al., 2003) or *Streptococci + C. albicans* dual-species biofilm growths (Dutton et al., 2016) and we wanted to determine if some of these genes are affected by GAs treatment. A semi-quantitative RT-PCR analysis was used to examine variation in the expression of genes related to biofilm formation like *cshA, ldh, gapdh, gftG1, scaA*, and *scaR* for *S. gordonii* and *CSH1, ZRT1, NRG1* and *PRA1*, for *C. albicans.* Treatment with GAs significantly reduced the expression of genes including *scaA, gapdh* and *gtfG1* in *S. gordonii* mono-biofilms whereas in dual biofilms, genes like *scaA, ldh* and *cshA* were reduced in their expression when compared with their respective controls (Figure 5A). Interestingly, the expression of *ldh* was enhanced 9-fold in GAs treated *S. gordonii* mono-biofilms but not in dual biofilms (Figure 5A). In mono-biofilms of *C. albicans*, the expression of *NRG1* was increased 2 fold in GAs treated samples compared to the untreated control (Figure 5B). No change was observed for *NRG1* and *CSH1* in GAs treated dual biofilms. The expression of *PRA1* was increased 2 fold in GAs exposed *C. abicans* mono-biofilms. Whereas, in dual-species biofilms the expression of *PRA1* was inverse in GAs treated biofilms. In contrast, *ZRT1*, the regulator of PRA1, was overexpressed about 5 fold in dual-species biofilms in the presence of GAs but not in the GAs exposed *C. albicans* mono-biofilms.

**Figure 5.**
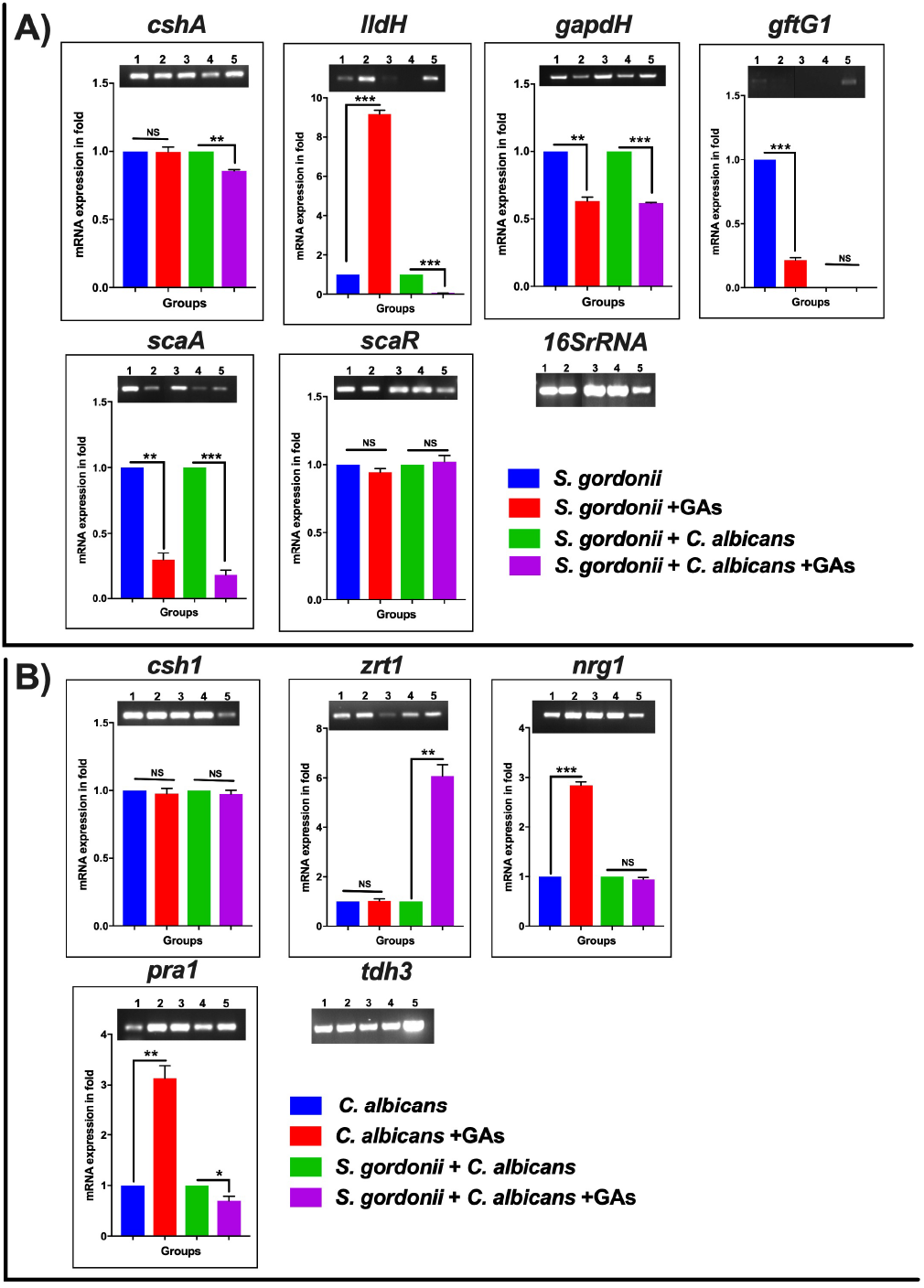
The mRNA expression level of biofilm genes as determined by RT-PCR. A) Representative semiquantitative mRNA expression profile for streptococcal primers showing the amplicons of mono- and dual-species biofilms. 1. *S. gordonii*, 2. *S. gordonii* + GAs, 3. *S. gordonii* + *C. albicans*, 4. *S. gordonii* + *C. albicans* + GAs, 5. Positive PCR control (gDNA used as template). Bar graphs represent the densitometry analysis of respective genes and a constant level of expression of 16S rRNA. B) Representative semiquantitative mRNA expression profile for candida primers showing the amplicons of mono- and dual-species biofilms. 1. *C. albicans, 2. C. albicans + GAs, 3. S. gordonii + C. albicans, 4. S. gordonii + C. albicans* + GAs, 5. Positive PCR control (gDNA as template). Bar graph represents the densitometry analysis of respective genes and a constant level of expression of *TDH3.* The results represent means ± standard deviations for three independent experiments. NS-not significant, \**p* <0.05, \*\**p*<0.01, \*\*\**p*< 0.001.

### GAs inhibit the GAPDH activity

To assess the potential inhibitory activity of GAs against the GAPDH from *S. gordonii*, we cloned the gene, overexpressed and purified the rGAPDH protein using the *E. coli* expression system (Figure 6A). The purified rGAPDH migrated at an apparent molecular weight of ~40 kDa and reacted to polyclonal anti-GAPDH antibody (Figure 6B). We next tested the effect of GAs (100 and 200μM) against the purified rGAPDH protein (0.1 μM). The assay depends on the conversion of glyceraldehyde-3-phosphate to 1,3-diphosphoglycerate by GAPDH enzyme in the presence of NAD. Interestingly, GAs appears to bind to GAPDH protein and block its enzyme activity in a dose-dependent manner. At 200μM concentration, GAs block the activity of GAPDH completely when compared to the reaction without GAs where it shows strong enzyme activity (Figure 6C).

**Figure 6.**
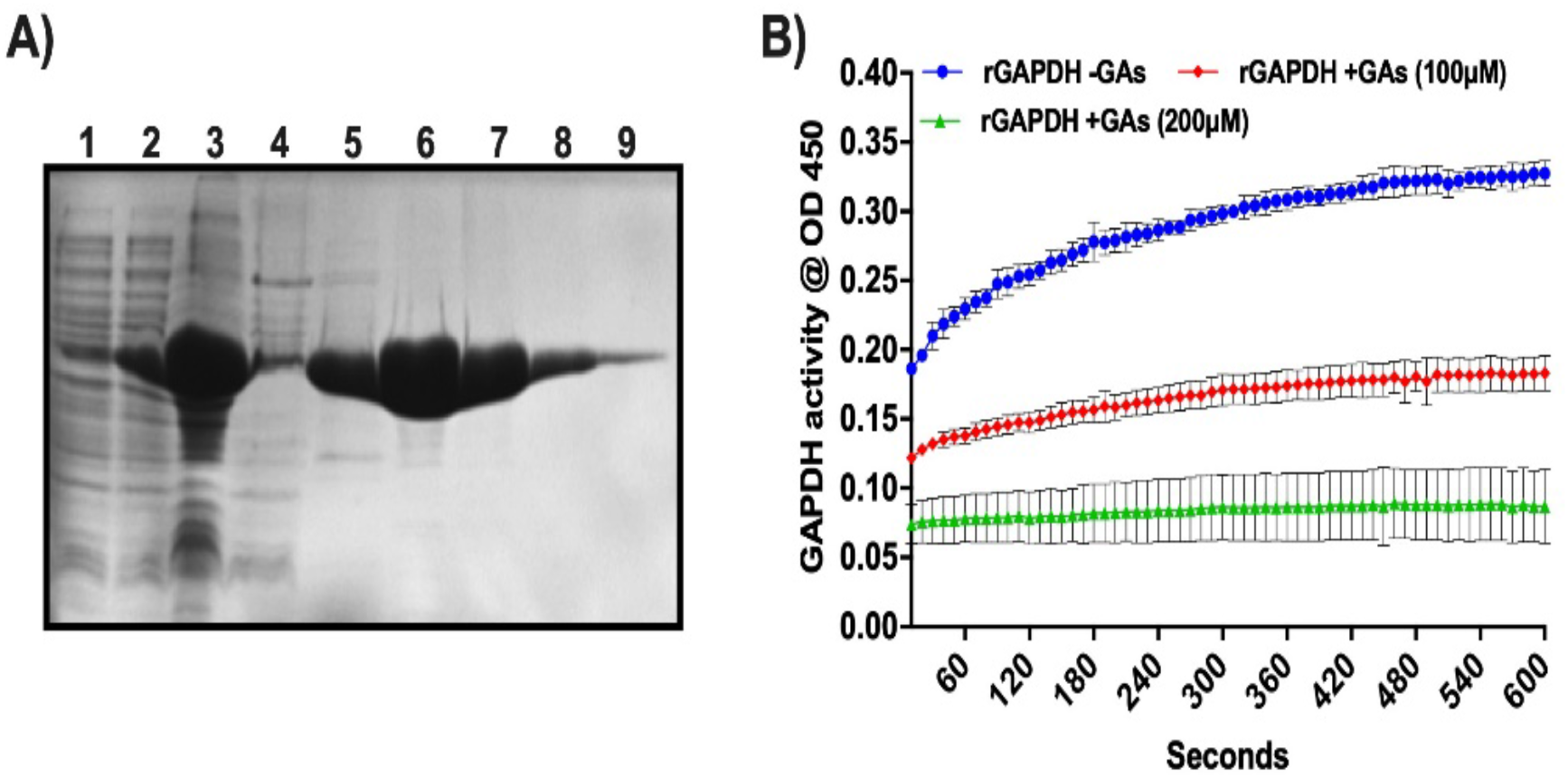
Purification of *S. gordonii* rGAPDH and determination of its enzyme activity. A) SDS-PAGE gel showing purified fractions of rGAPDH protein from *E. coli* cell lysates. 1. Uninduced whole cell lysate, 2. Induced whole cell lysate, 3. French pressed cell lysate, 4. Unbound fraction, 5 to 9 – 100 mM, 150 mM, 200 mM, 250 mM, 300 mM and 350 mM imidazole eluted fractions, respectively; B) Western blot of rGAPDH protein against anti-GAPDH antibody. Lanes, 1 & 2 – Purified rGAPDH protein at two different concentrations; Measurement of GAPDH activity in the presence and absence of GAs.

## Discussion

Microbial infection in the oral cavity of humans is biofilm-associated where a significant proportion of infection was mixed biofilms. *S. gordonii*, an early colonizer of the oral cavity forms an adhering biofilm on oral surfaces via cell surface adhesins (Bamford et al., 2009), leads to stable colonization in the oral cavity and also attaches to *C. albicans* hyphae via protein-protein interactions (Holmes et al., 1996). In addition, *S. gordonii* colonization on the tooth surface allows other microbes to adhere and develop mixed biofilms such as dental caries, which is the most prevalent human oral diseases, especially among the children. We have investigated the *S. gordonii* mono- and *S. gordonii – C. albicans* dual-species biofilms and their inhibition by gymnemic acids (GAs) *in vitro.* GAs, a medicinal plant-derived small molecule, was shown to prevent *C. albicans* yeast-to-hypha transition and hyphal growth without affecting its viability or yeast growth rate (Vediyappan et al., 2013). However, GAs’ effect on bacterial and or bacterial-fungal mixed biofilms are unknown. GAs are a family of triterpenoid saponin compounds which are the major active principles of *Gymnema sylvestre* plant leaves. The extract of this plant is widely used for its various medicinal properties including lowering blood glucose activity in diabetic patients and reducing obesity (Porchezhian and Dobriyal, 2003; Leach, 2007; Zuniga et al., 2017).

Antibiofilm efficacy of GAs was investigated in terms of CV staining and eDNA reduction, where the results found to be significant compared to untreated controls. This is the first report to provide evidence that the GAs shows antibiofilm efficacy against both mono- and dual-species biofilms of *S. gordonii* and *C. albicans.* The microbial biofilms are protected by self-produced exopolysaccharides (EPS). EPS are generally made up of different types of polysaccharides, proteins, glycoproteins, glycolipids, and eDNA. The importance of eDNA release during early stages of biofilm is to preserve the structural firmness, enhancing the mixed biofilm and protection against antimicrobial agents (Mulcahy et al., 2008;Jack et al., 2015;Jung et al., 2017). Therefore, reduction of eDNA accumulation and other components could substantively diminish the development of biofilm formation. As such, we found that GAs was able to reduce a significant amount of eDNA being released by both mono- and dual-species biofilms (Figures 1 & 2).

It was reported earlier that *S. gordonii* cells form surface fibrils which have multiple properties like cell surface hydrophobicity, co-aggregate with other oral bacteria, saliva-coated hydroxyapatite (sHA) and bind to host fibronectin (McNab et al., 1996;Back et al., 2017). These results emphasize that fibril-mediated attachment is the critical factor for the initial oral colonization for Streptococci. In the present study, we observed an extracellular nanofibrillar-mediated attachment of *S. gordonii* cells to sHA by SEM. Interestingly, these nanofibrils were not peritrichous as previously reported (McNab et al., 1999) and instead, the scattered fibrils were attached to neighboring streptococci cells, sHA substratum and to *C. albicans* hyphae (Figures 3 & 4A-D) confirming its role in adherence. To our surprise, synthesis of these fibrils were abolished in the GAs treated *S. gordonii* biofilms. These fibrils could be related to EPS and we believe GAs might be affecting their synthesis and or their incorporation into the biofilms. One of the unexpected findings of *S. gordonii-C. albicans* mixed biofilms was the formation of short fibrils from the *C. albicans* hyphae (Figure 4A & B). These fibrils show attachment to neighboring hypha and to the sHA substratum. This shows that there is an enhanced mutual synergism between these two microbes. However, in GAs treated mixed biofilms, these fibrils were absent (Figure 3L) and found significant inhibition of biofilms. Djaczenko and Cassone (1971) have reported the presence of fimbriae in *C. albicans* yeast cells and known to contain mannosylated glycoprotein (Yu et al., 1994). We believe the fibrils that we observe in hyphae could be different from the fimbriae described above. For example, the fimbriae reported by Djaczenko and Cassone (Djackenko and Cassone, 1971) were found on the surface of ‘yeast cells’ grown on agar plates for several days. These fimbriae are short and continuous throughout the cell surface of mother yeast cells but very little on the daughter cells.

In contrast, our results show the fibrils are discontinuous and found only from hyphae of *S. gordonii-C. albicans* co-cultured biofilms where they have close contacts with abiotic or biotic surfaces (Figure 4). These fibrils were not observed in biofilms grown in the presence of GA, suggesting that GA can prevent adhesive fibrils, in part, by inhibiting its synthesis and or hyphae associated mannoproteins.

To understand the mechanisms of biofilms inhibition by GAs, we next determined the expression of a few selected genes that have predicted roles in the growths of *S. gordonii* and *C. albicans* biofilms. EPSs are the core parts for the assembly and maintenance of biofilm architectural integrity in the oral cavity. The oral streptococci produce glucosyltransferase enzymes, Gtfs, that catalyze extracellular glucose into glucan polymer, which helps the streptococci adhere to the tooth surface and to the surfaces of other oral microbes. *S. gordonii*, the primary colonizer of the oral cavity, produces *gtfG* (Vickerman et al., 1997).

The RT-PCR analysis of *S. gordonii* biofilm cells shows basal level expression of *gtfG.* However, GAs treatment reduced its expression, signifies the inhibitory potential of biofilm glucan by GAs. This result agrees with SEM data where the *S. gordonii* biofilms treated with GAs show absence of adhesive fibrils when compared to the control biofilm where the fibrils can be seen between the biofilms cells and on the sHA (Figure 5). The other roles of Gtfs include glycosylation of adhesive proteins such as GspB of *S. gordonii* and Fap1 of *S. parasanguinis* (Zhu et al., 2015). GAs are known to bind several proteins including glucose transporter (Wang et al., 2014), taste receptors T1R2/T1R3 (Sanematsu et al., 2014), and Liver X-receptor (LXR) that regulates lipid metabolism in the liver (Renga et al., 2015). It has been reported that administration of GAs containing fraction, GS4, decreased the glycosylated hemoglobin (HbA1c) and glycosylated plasma protein in diabetic patients (Baskaran et al., 1990) and a similar mechanism may occur in microbial biofilms. Bacterial Gtfs play a critical role in enhancing the accumulation of *C. albicans* cells during mixed biofilms growths (Ellepola et al., 2017). Similarly, mannosyltransferase genes *(MNT1/2)* of *C. albicans* are involved in *O*-mannosylation of proteins in the hyphal surfaces that allow *S. gordonii-C. albicans* interaction and promote mixed biofilms (Dutton et al., 2014). In a separate study, we have observed that the treatment of *C. albicans* biofilms with GAs affected many of its mannosylated proteins (McMillan et. al. manuscript in preparation).

Thus, GAs appears to modulate glycan transferases from both *S. gordonii* and *C. albicans*. GAs may also affect the polysaccharide synthesis pathway in *S. gordonii* biofilms, through a reduced *gtfG* expression and or its enzyme activity. Further, Gtfs use metal co-factor Mn^2^+ for enzyme catalytic activity (Zhu et al., 2015) and the downregulation of *scaA*, the gene that encodes Mn^2^+ binding lipoprotein, in GAs treated *S. gordonii* mono-as well as dual-species biofilms (Figure 5) may also contribute to the reduction of adhesive fibrils/polysaccharides. For growth and survival in the human host, *S. gordonii* will have to acquire Mn^2^+ with the help of ScaA, a prominent surface antigen. It has been shown that inactivation of *scaA* gene resulted in both impaired growth of cells and >70% inhibition of Mn^2^+ uptake (Kolenbrander et al., 1998).

Oral bacteria including, *S. gordonii*, can sense the redox status of the biofilm niche and respond accordingly. Among the genes examined for differential expression in biofilms, we found lactate dehydrogenase *(ldh)* is one of the highly upregulated genes in GAs treated biofilms of *S. gordonii* (Figure 5). The ldh enzyme interconverts pyruvate into lactate and back, as it converts NADH to NAD and back. In GAs treated *S. gordonii*, ldh may be converting lactate into pyruvate as the *gapdh* mRNA is downregulated in GAs treated mono- or mixed biofilms of *S. gordonii* but not in *C. albicans.* GAPDH uses NAD during glycolytic activity and the reduced amount of GAPDH may lead to the accumulation of NAD, which in turn activates the overexpression of *ldh* through a redox-sensing system (Bitoun and Wen, 2016). To determine if GAs has any effect on GAPDH enzyme activity, we cloned the *gapdh* gene from *S. gordonii*, overexpressed in *E. coli* and tested the purified rGAPDH with or without GAs. We found the inhibition of rGAPDH enzyme activity in a dose-dependent manner (Figure 6). GA was shown to inhibit rabbit GAPDH enzyme activity (Izutani et al., 2005). Maeda et. al. have showed that oral streptococcal *(e.g. S. oralis, S. gordonii)* cell surface-associated GAPDH binds to the long fimbriae (FimA) of *Porphyromonas gingivalis* (Maeda et al., 2004a;Maeda et al., 2004b) and play a role in the development of oral polymicrobial biofilms (Kuboniwa et al., 2017). In addition to glycolytic function, GAPDH is also a moonlighting protein and known to carry out multiple functions (Sirover, 2017). It is worth mentioning that natural products (anacardic acid and curcumin) are shown to bind and inhibit *S. pyogenes* GAPDH activity, one of the major virulence factors (Gomez et al., 2019) and the GAPDH serves as a drug target in other pathogens (Freitas et al., 2009) as well. GAs appear to impact on *S. gordonii* GAPDH both at the transcriptional and translational level and could account, at least partially, for the observed inhibition of *S. gordonii* growth or biofilm. Comparison of amino acid sequences of both *S. gordonii* and *C. albicans* GAPDH revealed about 50% similarity and thus GAs impact on them could be different. In fact, the expression of GAPDH gene in *C. albicans (TDH3)* biofilms grown in the presence or absence of GAs is not affected (Figure 5B, *TDH3* RT-PCR bands). However, GAs impact on *C. albicans* GAPDH (Tdh3) enzyme activity and its role in biofilms can’t be ruled out and remains to be determined. Further, the global gene expression and biochemical analyses could reveal the complete mechanism(s) of GAs-mediated inhibition of *S. gordonii* mono- and mixed biofilms.

Among the genes examined in *C. albicans* mono- or dual-species biofilms, *NRG1, PRA1* and *ZRT1* are the most differentially expressed genes. It is well known from the literature that Nrg1 of *C. albicans*, is a DNA binding protein that represses its filamentous growth (Braun et al., 2001). GAs treatment shows a significant increase of *NRG1* mRNA expression in *C. albicans* biofilms compared to control biofilms (Figure 5B), which may correspond to the observed yeast or pseudohyphal growth forms of *C. albicans* mono-species biofilm (Figure 3). However, no change of *NRG1* expression level was observed in dual-species biofilms yet their biofilms growths were inhibited underscoring the unknown regulatory mechanism in the GAs treated dual biofilms. *C. albicans* sequesters environmental zinc through a secreted protein, the pH-regulated antigen 1 (Pra1) and transports it through the membrane transporter (Zrt1) for its invasive growth in the host (Citiulo et al., 2012). *C. albicans* has biphasic mechanisms for its environmental and cellular zinc homeostasis and Pra1 expresses when cells are at pH 7 and above or at zinc limitation (Crawford et al., 2018;Wilson, 2019). GAs treatment to *C. albicans* mono-biofilm appears to cause zinc limitation and or change in cellular pH, which could be altered when grown with *S. gordonii* as mixed-species biofilms (Figure 5B).

Our understanding about mixed species biofilms in caries pathogenesis was not studied well and still in its infancy (Metwalli et al., 2013). It was well known that from various host defense factors, microbes in mixed biofilms act synergistically for their survival (Morales and Hogan, 2010;Xu et al., 2014a). Great attention is needed on mitis group streptococci *(Streptococcus gordonii, Streptococcus oralis, Streptococcus mitis, Streptococcus parasanguinis*, and *Streptococcus sanguinis*), where it forms multispecies biofilms when aggregating with other bacterial and fungal species (Xu et al., 2014b). These oral microbial infections pose a significant threat to public health as many pathogenic bacteria readily develop resistance to multiple antibiotics and form biofilms with additional protection from antibiotic treatment (Lebeaux et al., 2014). Currently available antimicrobial agents were most effective at drastically reducing the cell viability, rather than reducing the virulence via inhibiting the biofilm growth. For instance, fluoride is one of the proved agents for caries prophylaxis, however, excess use of fluoride causes fluorosis and hardening of cartilage.

Also, these synthetic antimicrobial agents lead to negative effects in the gastrointestinal system and several other side effects. We are in need of efficient antimicrobial agent which inhibit the biofilm formation, alike it should not exert selective pressure over oral microbiome. Recently, many studies have been targeted over medicinal plants in finding effective anticaries agents (Islam et al., 2008;Yang et al., 2017;Gartika et al., 2018;Henley-Smith et al., 2018). Medicinal plants have been used to prevent and treat microbial diseases since ancient times, which can target several antigens or pathways of the pathogens for inhibition without adverse effects. Earlier studies on some medicinal plant extracts display biofilm inhibition through hindering hydrophobic property of *S. mutans* (Nostro et al., 2004;Khan et al., 2012). Any antimicrobial agent that reduces or hinder these types of interactions/attachment will be a novel strategy to overcome oral infection. Interestingly, our GAs treatment shows a significant reduction in both mono- and dual-species biofilms and appear to act via more than one mechanisms. GAs affect the transcription of *S. gordonii gapdh* and its enzyme activity in addition to *gtfG1* which is involved in glucan polysaccharide synthesis. Further, GAs can able to curtail the development of nanofibrils both from *S. gordonii* and *C. albicans* that mediate cell-cell and substrate adhesion. In summary, our findings offer an anti-virulence approach for preventing mixed oral biofilms and by further optimization, the natural products could be a useful source for developing mixed biofilm inhibitors.

## Acknowledgements

The authors acknowledge the generous support of Streptococcal strains by Indranil Biswas, KUMC, Kansas City. We thank American Heart Association for SDG grant, Kansas Idea Network of Biomedical Research Excellence (K-INBRE) CORE Facility and Johnson Cancer Research Center, KSU for IRA funding supports to GV and K-INBRE postdoctoral support to RV. We also thank Erick Saenz-Gardea’s assistance under the Developing Scholar Program during the initial phase of this project. We acknowledge the SEM technical support by MAI, KU. This project was supported by an Institutional Development Award (IDeA) from the National Institute of General Medical Sciences of the National Institutes of Health under grant number P20 GM103418. The content is solely the responsibility of the authors and does not necessarily represent the official views of the National Institute of General Medical Sciences or the National Institutes of Health.

## Author Contributions

G. V. designed the study. R.V and G.V. conducted the experiments, analyzed the data and wrote the manuscript.

## Conflict of Interest Statement

We have no conflicts of interest to disclose.

